# Physical mechanisms of ultrasonic neurostimulation of the retina

**DOI:** 10.1101/231449

**Authors:** Mike D. Menz, Patrick Ye, Kamyar Firouzi, Kim Butts Pauly, Butrus T. Khuri-Yakub, Stephen A. Baccus

## Abstract

Focused ultrasound has been shown to be effective at stimulating neurons in vivo, ex vivo and in vitro preparations. Ultrasonic neuromodulation is the only non-invasive method of stimulation that could reach deep in the brain with high spatial-temporal resolution, and thus has potential for use in clinical applications and basic studies of the nervous system. Understanding the physical mechanism by which energy in a high acoustic frequency wave is delivered to stimulate neurons will be important to optimize this technology. Two primary candidates for a physical mechanism are radiation force, the delivery of momentum by the acoustic wave, and cavitation, oscillating gas bubbles. We imaged the isolated salamander retina during ultrasonic stimuli that drive ganglion cell activity and observed micron scale displacements consistent with radiation force. We recorded ganglion cell spiking activity with a planar multielectrode array and changed the acoustic carrier frequency across a broad range (0.5 - 43 MHz), finding that increased stimulation occurs at higher acoustic frequencies, a result that is consistent with radiation force but not cavitation. A quantitative radiation force model can explain retinal responses, and could potentially explain previous in vivo results in the mouse, suggesting a new hypothesis to be tested in vivo. Finally, we found that neural activity was strongly modulated by the distance between the transducer and the electrode array showing the influence of standing waves on the response. We conclude that radiation force is the physical mechanism underlying ultrasonic neurostimulation in the ex vivo retina, and that the control of standing waves is a new potential method to modulate these effects.

## Introduction

Ultrasonic neuromodulation has been demonstrated in many different experimental preparations including human (Lee et al., 2015; 2016a; Legon et al., 2014; Monti et al., 2016), monkey (Deffieux et al., 2013), sheep (Lee et al., 2016b), rat (Younan et al., 2013), mouse (Kamimura et al., 2015; Ye et al., 2016; King et al., 2014; King et al., 2012; Tufail et al., 2010; Tyler et al., 2008), and ex vivo salamander retina (Menz et al., 2013). The capability of ultrasound to reach any brain structure noninvasively through the skull, and the highly developed technology to deliver ultrasound make this approach promising as a potential method for basic studies of neural function and clinical applications. Yet results in different preparations have varied, including both excitatory and inhibitory effects. The development of this approach would benefit greatly from a quantitative understanding of the mechanisms of ultrasonic neuromodulation, allowing the process to be optimized in terms of efficacy of stimuli, efficiency and spatiotemporal distribution of effects.

In the process of transduction of a stimulus into a biological response, one can distinguish the physical mechanism such as acoustic pressure or thermal energy from the biophysical mechanism that senses that energy, including changes in membrane capacitance or particular ionic channels. Here we focus on the physical mechanism by which an acoustic wave is converted into an effective stimulus for a neuron, a process that is currently not understood. The leading candidates for physical mechanism are radiation pressure, the process by which an absorbed or reflected wave delivers momentum, and cavitation, which includes the stable or unstable formation of bubbles, creating a mechanical disturbance, and thermal energy.

Radiation force is a nonlinear effect proportional to the intensity of the acoustic wave, thus creating a continuous, non-oscillating force for a stimulus of constant amplitude (Rudenko et al., 1996). By this mechanism a carrier wave with a frequency too high to have a direct biological effect can be converted into a low frequency mechanical force with dynamics of the envelope of the wave. When radiation force is exerted on a liquid, this results in bulk flow of fluid known as acoustic streaming. Tissue attenuation increases with carrier frequency, therefore radiation force will increase with frequency, as will heating due to absorption.

Cavitation can occur if the acoustic pressure wave becomes sufficiently negative, causing gas bubbles to form that oscillate at the carrier frequency synchronously with the changing acoustic pressure (Nightingale et al., 2015). Inertial cavitation occurs when those oscillations change in size and eventually burst the bubble, creating a destructive violent event. In stable cavitation the bubble does not burst, and is hypothesized to produce safe neuromodulation. Cavitation is less likely at higher carrier frequencies because it becomes more difficult to sustain oscillations in the bubble.

Here we use optical imaging to measure displacements in the retina, and vary the acoustic frequency to test which of these mechanisms is most likely. We find that ultrasonic stimulation in the retina is consistent with a model whereby radiation force produced micron-scale mechanical displacements. The acoustic frequency dependence is consistent with radiation force but inconsistent with cavitation. In addition, we see that standing waves influence the effects of ultrasound. We conclude that radiation force is the primary physical mechanism for ultrasound to stimulate the retina.

## Results

### Radiation force causes physical displacement within the retina

We imaged the retina with a two-photon laser-scanning microscope after applying the membrane dye FM4-64 to the bathing medium (Fig. 1). We recorded a stack from the MEA up to the photoreceptor level while repeatedly stimulating the retina with ultrasound (43 MHz carrier, one second on, one second off, I_SP_ = 40 W/cm^2^). Since ultrasound stimulation and the scanning laser were uncorrelated, all points in the volume were imaged relative to the onset of the ultrasound stimulus, though on different trials. Using the time that the laser scanned each pixel, we reconstructed a movie of the average intensity at each pixel for the entire volume at a 10 ms resolution with respect to the ultrasound stimulus (see Supplemental Movie 1). At the onset of ultrasound, a sudden displacement was observed that lasted the duration of the stimulus. This displacement was centered at the ultrasound focus, was greatest in the outer retina, and decreased to zero near the ganglion cell layer. Lateral to the focal point, the displacement became progressively more lateral. In addition, there was relatively little movement in the ganglion cell layer on top of the MEA.

**Figure 1.**
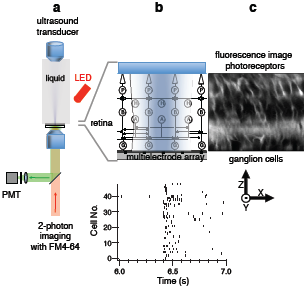
Experimental configuration for ultrasonic stimulation of the retina ex vivo. **a**, Schematic of ultrasound transducer mounted vertically and immersed in perfusion fluid with the focal point on the retina. Two-photon imaging is performed from below while a red LED from above can be used for visual stimulation. **b,** Top, An expanded view showing that the retina placed ganglion side down on an MEA. The ultrasound field (shading) spans the width of the entire retina. The actual ultrasound field is shown in (Supplemental Figure 1) Bottom, A population of ganglion cell spiking activity recorded with an MEA in response to ultrasound. **c**. Retinal image using the dye FM4-64 (showing cell membranes and processes) is a slice in the XZ plane.

These displacements between steady-state ‘ON’ and ‘OFF’ ultrasound were converted into a vector field of displacements using image processing software (see Methods) (Fig. 2a). Anatomically, the inner plexiform layer (IPL, Fig 2a) is a large network of entangled small processes and likely has different mechanical properties than the cell body layers above and below. From the vector field we observed that displacement below the IPL and above the MEA was very small. Other fluctuations in displacements were observed that could be a consequence of inhomogeneity in the retina. Confirming observations from the movie, there was little displacement in the ganglion cell layer.

**Figure 2.**
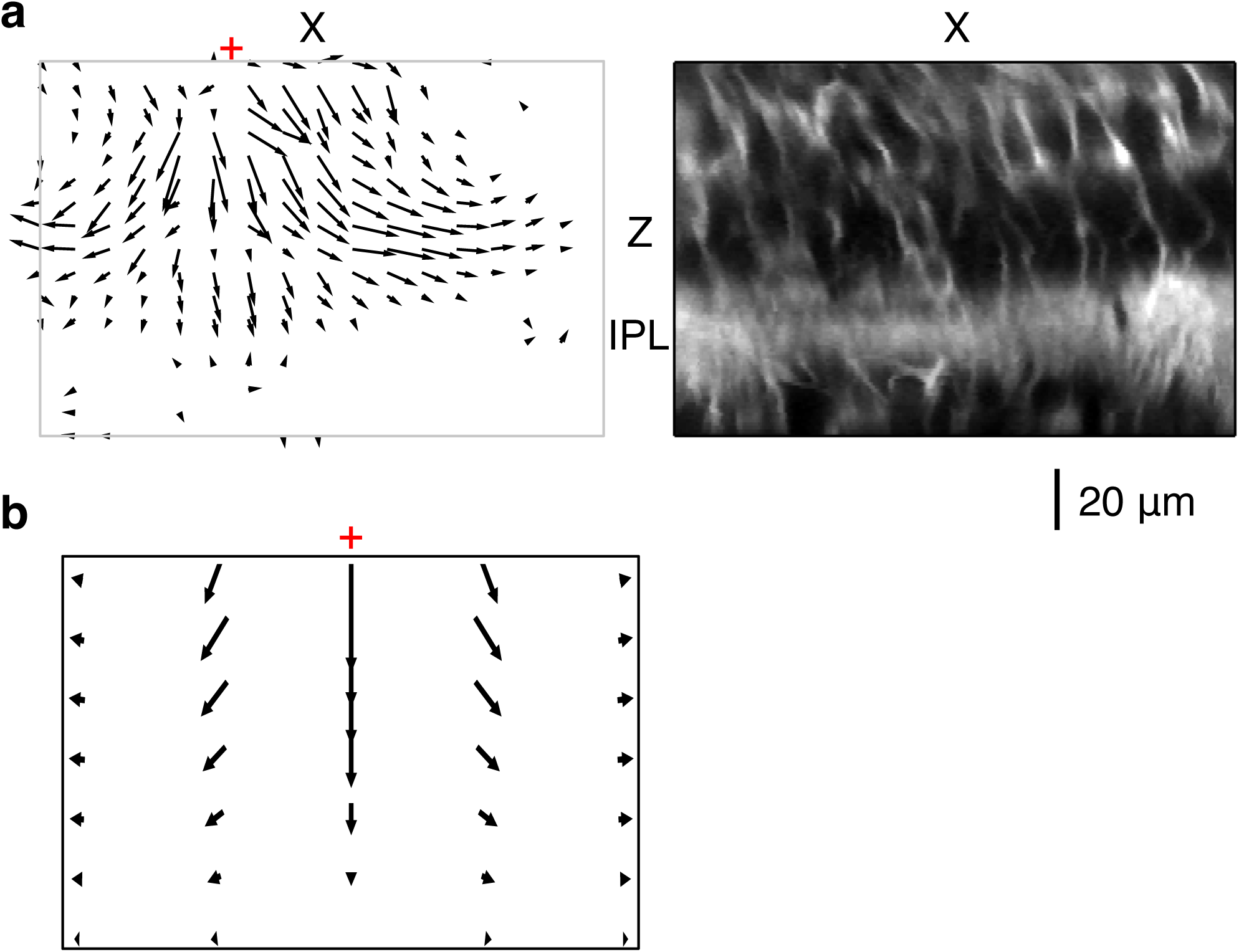
Ultrasonic radiation force causes displacement in the retina. **a**, Left, At 43 MHz and 40 W/cm^2^ I_SP_, an XZ slice through the retina near the focus of a vector field showing displacement (relative magnitude and direction). The vector field was computed from the image at steady ultrasound ON relative to ultrasound OFF. Red cross indicates the center of focus. Right, XZ slice with ultrasound OFF used in **a**. Left. **b**, Vector field of displacement using a simulation of radiation force acting on the retina, E (Young’s modulus of elasticity) = 2.4 kPa, maximum displacement is at center top (4 μm). Scale of vectors is different than the scale of the image. Red cross is center of focus.

To interpret the potential mechanism of this displacement, we modeled the expected mechanical response of the retina from radiation force using finite element analysis (COMSOL). A key parameter for this calculation is the Young’s modulus of elasticity for the retina. However, the literature has values that vary three orders of magnitude, depending on the method of measurement (McKee et al., 2011). We thus allowed the Young’s modulus to be a free parameter and fit the model to account for the maximum observed displacement, which was 4 microns. The resulting value of Young’s modulus was 2.3 kPa, which is close to the range found in the inner retina (0.94-1.8 kPa) with the scanning force microscopy method (Franze et al., 2011). Our displacement results indicate the outer retina is more relevant, and Young’s modulus could be somewhat higher in the outer retina. The general features of the model simulation vector field qualitatively match the experimental vector field of displacement: large downward motion in the outer retina right under the focus which decreases to zero at the level of the MEA (Fig. 2b). Lateral to the focal point the displacement was more lateral and less downward. In the simulation, the retina was modelled as a homogeneous medium, so features such as the large change in displacement at the boundary of the IPL were not captured.

To quantify the temporal dynamics of displacement, we found a region with high local contrast that had the largest displacement, and examined the change of displacement in 10 ms time bins (Fig. 3a). There was 4 microns of vertical displacement which occurred very rapidly (< 10 ms) consistent with radiation force (Prieto et al., 2013). The fast onset of displacement is consistent with the fast response of neurons to ultrasonic stimulation (Menz et al., 2013). The recovery to baseline was slower and was fit by double exponential with time constants of 21 ms and 304 ms (Fig 2b). This recovery reflected the elastic properties of the retina. In a typical experiment we use a dialysis membrane to hold the retina against the MEA. In the imaging data shown in (Fig. 1-3) a large hole was cut in the dialysis membrane so there was only fluid between the transducer and retina. This simplified the simulation of retinal displacement by radiation force shown in Fig 2b because the dialysis membrane has unknown mechanical and acoustic properties. We also performed this imaging experiment with the dialysis membrane in place (results not shown) and found similar qualitative result, although the maximum amount of displacement was only 2 microns. It appears that the membrane attenuates the ultrasound wave so that intensity and displacement is reduced in the retina. We can conclude that the experimentally observed displacement is caused by radiation force at power levels known to produce ultrasonic neurostimulation.

**Figure 3.**
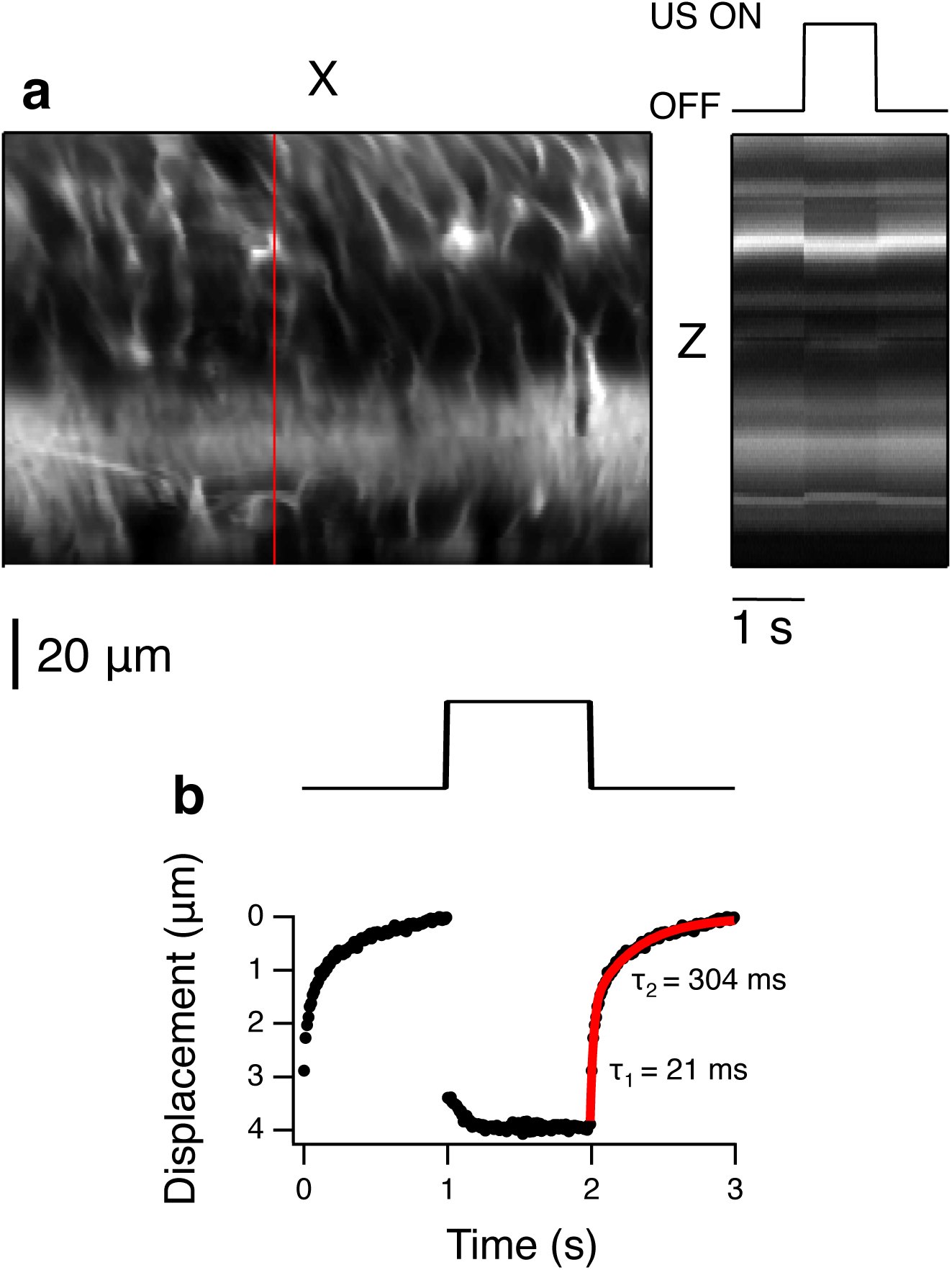
Ultrasound causes fast (< 10 ms) micron-scale tissue displacement. **a**. Left, XZ image slice, red line indicates the spatial cross section which is then shown as a function of time at right. Right, temporal changes during one second ultrasound OFF, one second ON, and one second OFF. **b**. A Gaussian was fit to the bright spot in each 10 ms time bin and the mean position plotted as a function of time. Stimulus trace showing the timing of ultrasound onset and offset. Bottom, Vertical displacement as a function of time. The relaxation after stimulus offset is shown fit with a double exponential (red curve).

### Relationship between displacement and ultrasonic neurostimulation

To examine the relationship between displacement and neural activity, we then compared measurements of these two quantities as a function of stimulus intensity. We imaged a level in the retina above the IPL midway through the retina, a level that showed considerable lateral displacement. We varied the ultrasound intensity from below the threshold of neural activation to above the level of a saturating response (Fig. 4). We computed the displacement at this level as a function of ultrasound intensity, and compare this relationship to that of the normalized firing as a function of ultrasound intensity taken from a different preparation (Fig. 4b). Neural activity was observed at a threshold of ~ 1 W/cm^2^, a level at the threshold of detectability of tissue displacement. The shapes of the two curves were different, with displacement increasing approximately linearly with stimulus intensity, and neural activity having a saturating dependence on intensity that was sigmoidal on a logarithmic scale. For each intensity value we plotted the normalized firing rate vs. displacement (Fig 4c). There was a rapid increase in firing over submicron values of displacement, after which neural activity saturated. This indicates that submicron scale displacement were correlated with neural activity, and gives a scale for the biophysical mechanisms that could transduce these displacements to produce activity.

**Figure 4.**
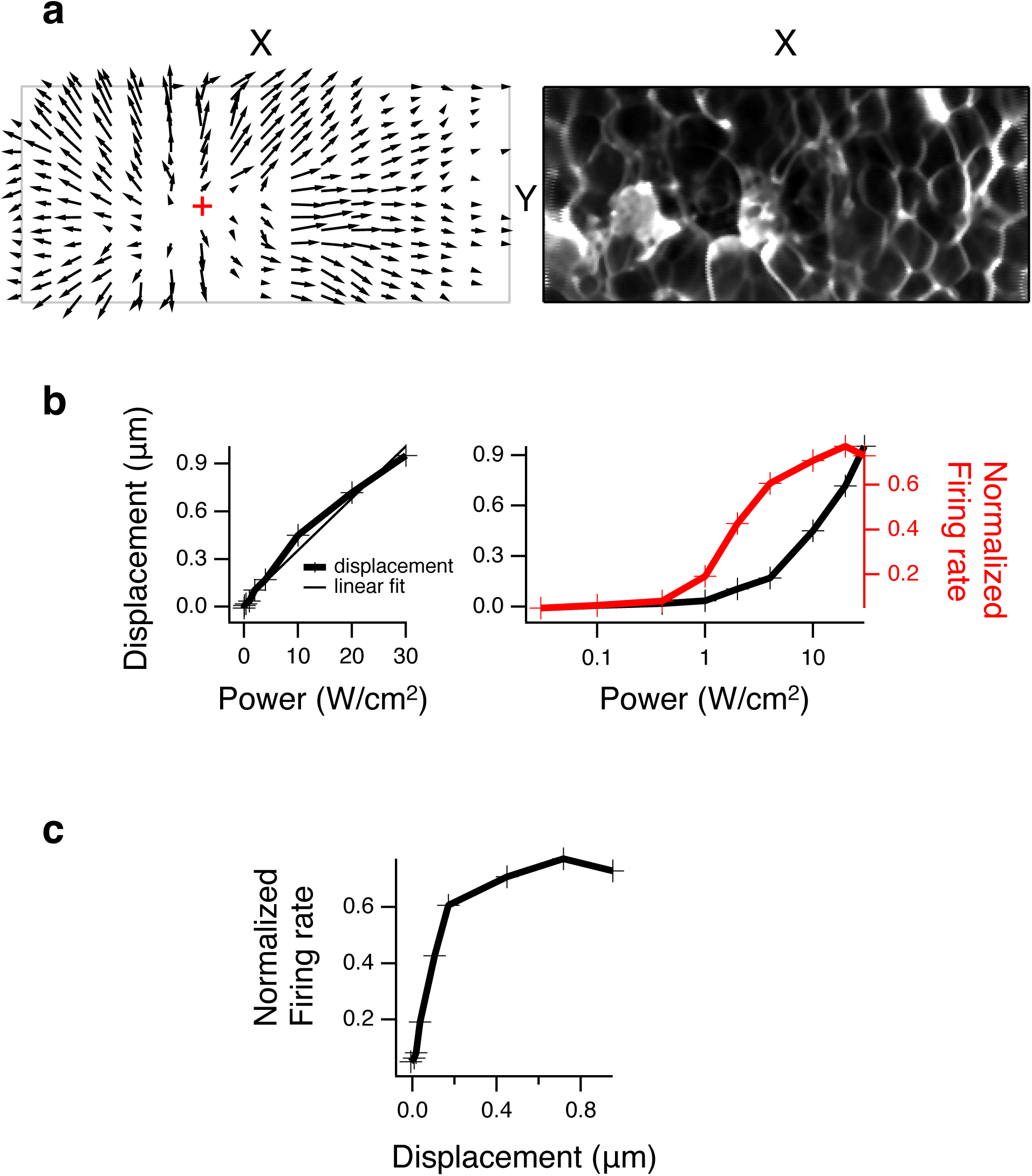
Relationship between displacement and neural activity. **a**, Left, An XY slice through the focus (red cross, point of minimal displacement) at a depth of maximal lateral displacement about midway through the retina that shows lateral displacement vector field in all directions moving away from the focal point. Right, XY slice with ultrasound OFF used in **a. b**, Left, Maximum lateral displacement (in XY for a given depth) is plotted vs. intensity. Data points are shown as “+”, thin line is a linear regression fit. Right, Normalized population firing rate is plotted vs. intensity and superimposed over the plot to the left, where the x-axis is now on a log scale. **c**, Normalized population firing rate is plotted vs. displacement from **b** for each intensity value.

### Relationship of response and acoustic frequency

Absorption increases with higher acoustic frequency, and thus both radiation force and heating are expected to increase with higher carrier frequency. In contrast, the probability of cavitation decreases with higher carrier frequency because of the shorter time interval available to cause a bubble to form out of solution and to keep it oscillating. Many protocols of ultrasonic neurostimulation use lower frequencies (~1 MHz) to allow sufficient energy to penetrate the skull, and it is conceivable that at lower frequencies a different mechanism such as cavitation is involved (Plaksin et al., 2016). We therefore changed carrier frequency in several steps between 43 MHz and 0.5 MHz to measure the effects of ultrasound at different frequencies on the retina.

To more completely characterize response at a given frequency we varied both pulse intensity and duration across a wide range for the 43 MHz transducer (Fig. 5a, Left). The pulse duration that generates a response at the lowest intensity is 100 ms. Pulse durations longer than this are useful to distinguish the separate effects of stimulus onset and offset, but there was no increase in response sensitivity with increasing duration (Fig. 5b). However, as pulse duration was decreased below 100 ms, greater intensity is required to achieve stimulation. This relationship is consistent with the threshold being proportional to the integration of the pulse to obtain total energy, as is also found in electrical stimulation (Boinagrov et al., 2014).

**Figure 5.**
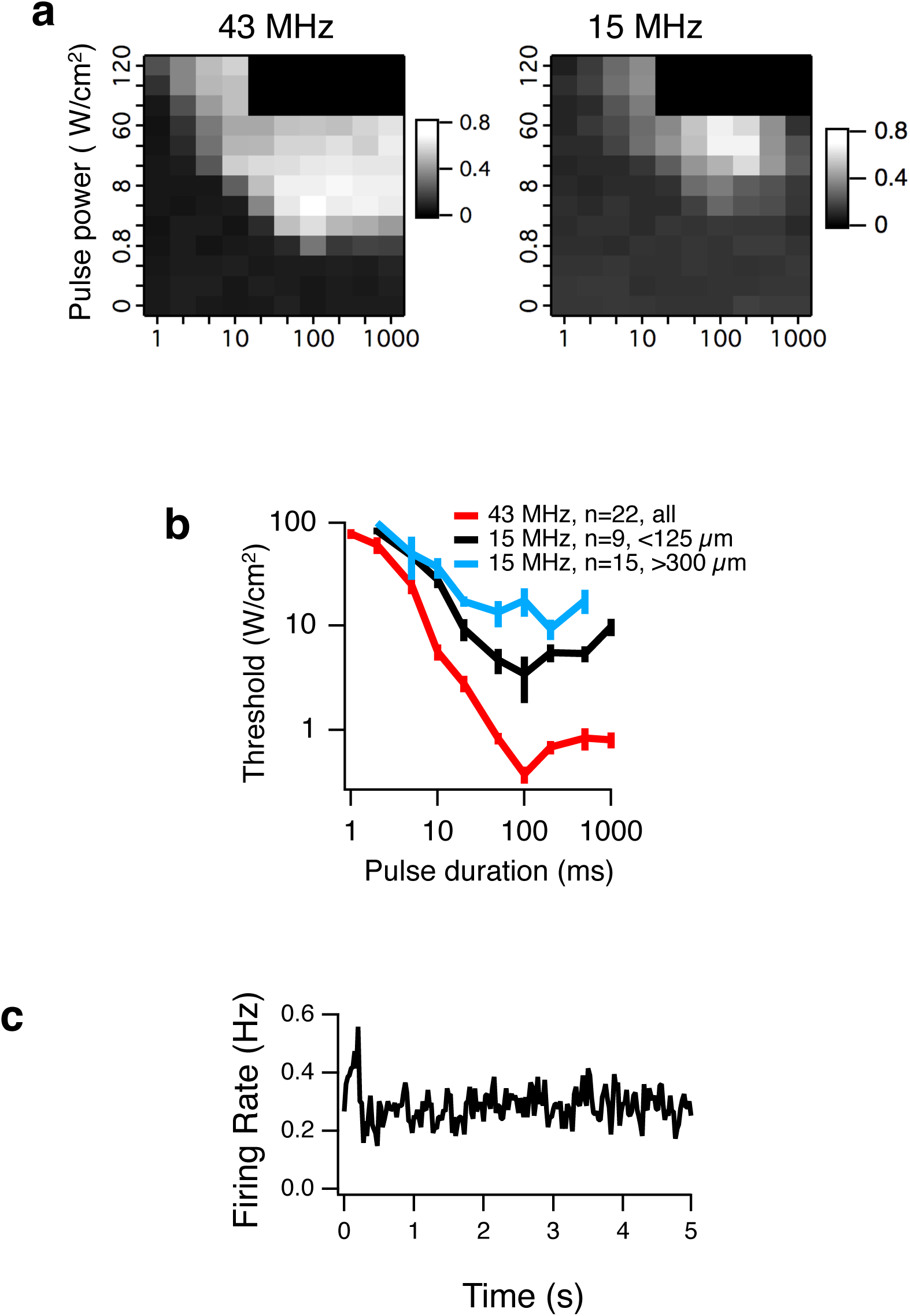
Ultrasonic stimulation at higher acoustic frequency has a lower threshold. **a**, Left, Normalized population firing rate is plotted as a function of intensity I_SP_ and pulse duration for 43 MHz. Right, Same plot for 15 MHz **b**, Threshold of stimulation as a function of pulse duration. Average thresholds across cells, (error bars are SEM) at each pulse duration for the three conditions, 43 MHz, 15 MHz for cells less than 125 μm from the focus, and 15 MHz for cells more than 300 μm from the focus. **c**, The population PSTH generates a very weak response to ultrasound (100 ms ON starting at time zero, repeated every 5 seconds) for a 500 kHz transducer at with I_SPPA_ = 1.6 W/cm^2^.

The same stimuli were used with a 15 MHz transducer. Because the focal area is larger with 15 MHz we also used an array with greater spacing between the electrodes (100 μm spacing in an 8x8 grid as opposed to 30 μm spacing in a 5x6 grid for the 43 MHz transducer). At 15 MHz a greater intensity was required to stimulate neurons compared to 43 MHz (Fig. 5a). At 15 MHz the 100 ms pulse duration was still optimal and this value is used as the default pulse duration in other experiments. Given the larger array used for 15 MHz we could segregate neurons into those closer to the focus and neurons that are farther away. Neurons that are closer to the ultrasound focus will experience higher radiation force and they have lower thresholds compared to neurons further away (Fig 5a, b). The higher thresholds at 15 MHz compared to 43 MHz is consistent with radiation force as the physical mechanism.

We then lowered the carrier frequency to be within the range used for in vivo ultrasonic neuromodulation (0.5-2 MHz). At these low frequencies attenuation is much lower, especially from the skull, allowing for transcranial non-invasive stimulation. Since attenuation is much less one would expect that if radiation force was the physical mechanism, much greater intensity would be required compared to 15 and 43 MHz to achieve stimulation. Although the response curve at 1.9 MHz was a similar shape to that of 15 and 43 MHz, it was shifted substantially to a higher threshold intensity, consistent with radiation force (Fig. 6b). Finally, we attempted stimulation at 500 KHz. However, as frequency decreases the focal volume increases, thus limiting the maximum peak intensity achievable at low frequencies. At 500 kHz, at the maximum achievable spatial peak power with our transducer (I_SP_ = 1.6 W/cm^2^) response of single cells to this stimulus could not be detected with significance, and was only detectable when averaging across a population of neurons (Fig 5c). The increase in neural activity with increasing acoustic frequency was qualitatively consistent with radiation force, and inconsistent with cavitation as a mechanism.

**Figure 6.**
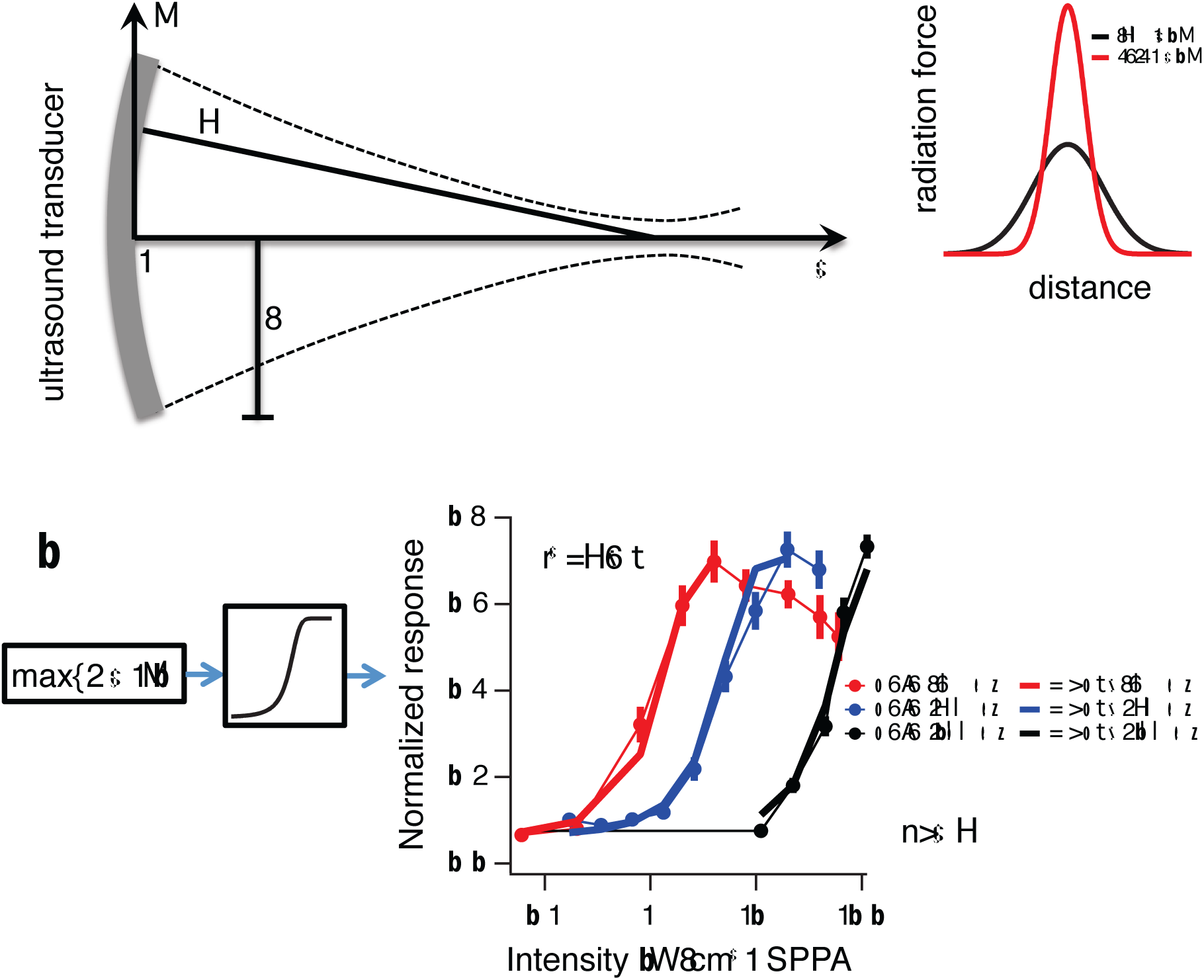
Radiation force model predicts retinal response across a range of acoustic frequencies. **a**, Left, An analytic expression was used to calculate radiation force in a cylindrical coordinate system based on transducer characteristics (*a* = radius of transducer, *d* = focal distance, *f*=frequency, *I* = intensity, *x* = axial distance, *r* = radial distance) (Rudenko et al., 1996, see methods). Right, A cartoon model of radiation force distributed over 1-D space for low (black) and high (red) frequencies. ***b***, Left, The maximum radiation force calculated from the above analytic expression is passed through an optimized sigmoidal non-linearity to generate the model responses.. Right, Normalized population response for 43, 15 and 1.9 MHz as a function of intensity, compared to the radiation force model output.

### Modelling radiation force to explain neural activity

We then tested whether neural activity could be fit with a single quantitative model of radiation force across the range of frequencies tested. We used an analytical model of radiation force valid for linear low-amplitude ultrasound in free space (Eqs 13,14,16 in (Rudenko et al., 1996), which has absorption coefficient, the carrier frequency, intensity, radius of the transducer, and focal length as parameters, to estimate radiation force in three-dimensional space expressed in a cylindrical coordinate system. From this model, we computed the maximum radiation force for each intensity and transducer, and then passed this value through a sigmoidal function to predict neural activity. The only free parameters defined this sigmoid, which was fixed across all acoustic frequencies. This model showed that the analytically computed maximum radiation force could be used to predict the neural response from 1.9 to 43 MHz with a single sigmoidal neural activation function (Fig 6). In summary, all of our results are consistent with radiation force as the physical mechanism of ultrasonic neurostimulation, including the expected quantitative dependence on acoustic frequency.

### Standing waves

Below the tissue, the glass surface of the MEA creates a large acoustic impedance mismatch, which we expect to reflect ultrasound. Thus the space between the transducer and MEA may form a cavity that could generate a standing wave, where locations spaced at one-half the acoustic wavelength, λ would experience destructive interference (nodes), and intervening locations experiencing constructive interference (anti-nodes). The relationship between standing waves and radiation pressure is well known in microfluidics (Bruus, 2012; Lenshof et al., 2012), where it is used to physically manipulate small particles, including individual cells. Radiation pressure is greatest at anti-nodes and smallest at nodes, causing tissue at nodes to be compressed by adjacent high pressure anti-nodes. Such mechanical pressure on tissue could have an additional influence on neural activity. We tested the neural effects of standing waves by simply changing the distance between the transducer and the MEA. This will not change the locations of the nodes and anti-nodes as they are fixed by the carrier frequency, but the change in cavity length will affect the amplitude of standing waves, with a maximal standing wave amplitude when the cavity length is a multiple of λ/2. Since the acoustic impedance of the MEA is much greater than tissue, the reflection will be accomplished with no change in phase. This means that there is an anti-node (high pressure) right at the border of tissue and MEA and a node (low pressure) located λ/4 away.

We chose to test the effects of standing waves at relatively low frequencies, 2.9 MHz (λ=517 μm, λ/4=129 μm) and 1.9 MHz (λ=789 μm, λ/4=197 μm), close to where most studies are conducted, yet high enough that we can still get robust responses, and the λ/4 distance is large and comparable to the thickness of the retina (~100 μm). The ultrasound stimulus was a continuous wave 100 ms pulse, which we had previously found to be optimal at higher frequencies (Fig 5b) and is very close to the 80 ms continuous wave pulse used for in vivo mouse stimulation (Ye et al., 2016). The stimulus was repeated every 5 seconds to minimize potential adaptation effects. A single 2.25 MHz transducer with relatively wide bandwidth was used for both frequencies, and intensities were measured by hydrophone separately at each frequency in free space.

We found that the firing rate of some cells was very strongly modulated by the distance between the transducer and the MEA with a period of λ/2, consistent with standing waves (Fig 7a,b). Across the population, we quantified the standing wave effect by computing the Fourier transform of the firing rate as a function of transducer distance, and examining the amplitude at a frequency of 2 cycles/λ as well as the phase angle of the response (Fig. 7c, at 2.9 MHz) relative to the starting position (0°, vertically mounted transducer) with the focus at the MEA and moving away from the MEA. The population showed that the response was modulated at a period of λ/2, consistent with a strong standing wave effect. We then tested whether standing waves were necessary for neurostimulation by tilting the transducer at an angle of 27° to vertical. Although a spatial interference pattern would still occur between the incident and reflected waves, the depth of modulation will not be as great as when the transducer is positioned vertically, and such a pattern would move with distance between the transducer and glass. We found that the tilted transducer condition still generated a response (Fig 7a), but the response modulation with distance was greatly reduced. At an angle of 27°, the average across the population showed a depth of modulation of 26 times less than when the transducer was vertical. We further tested that standing waves were also observed at 1.9MHz, λ/4=197 μm, using the same transducer, and similarly found that the population response was modulated at a period of λ/2 and that this average effect on the population diminished when the transducer was tilted an angle of 21° (Fig. 7d). From these results we conclude that standing waves influence the response, but are not necessary for stimulation.

**Figure 7.**
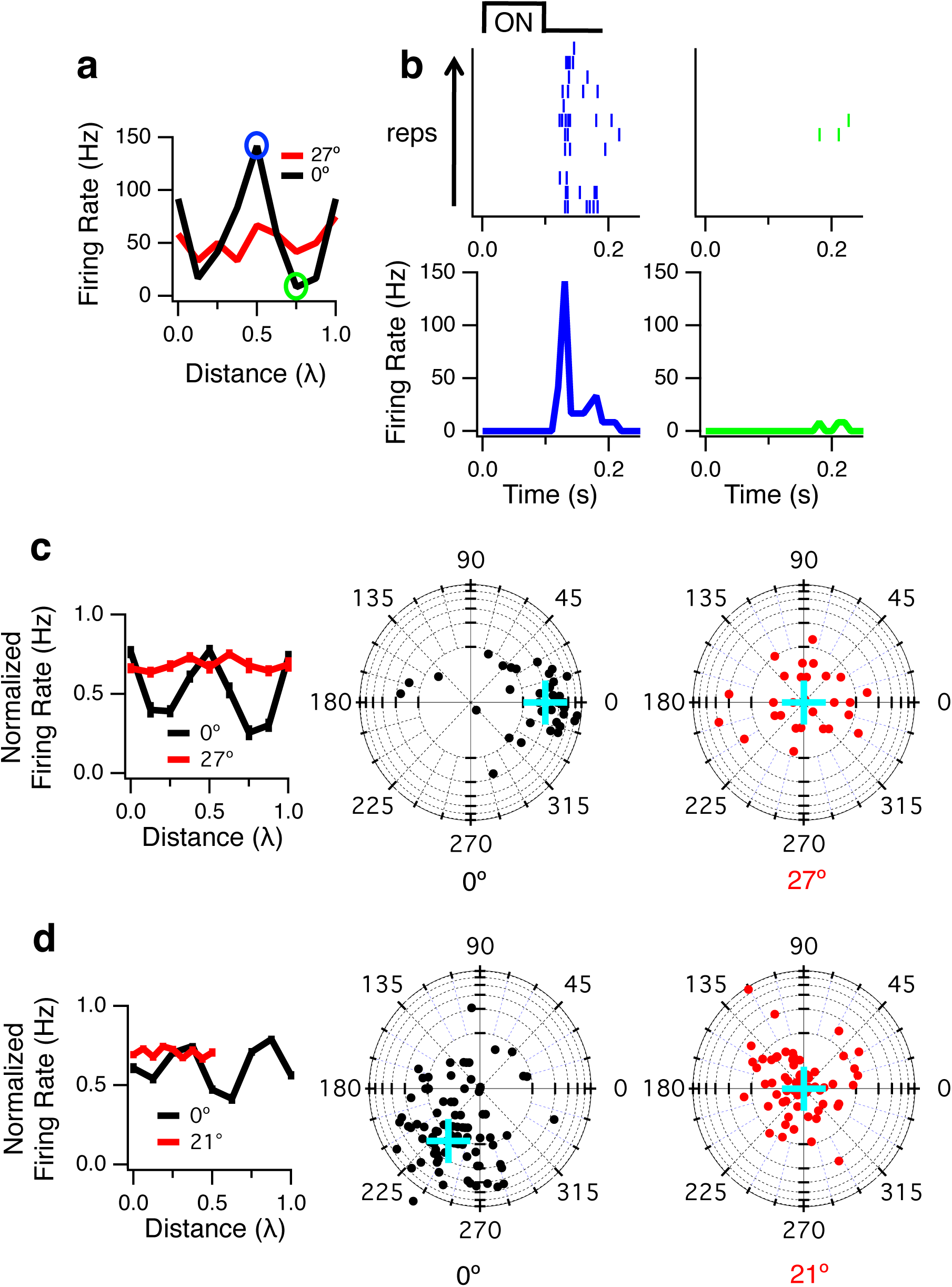
Standing waves modulate ultrasonic neurostimulation. A 2.25 MHz transducer was operated with a carrier frequency of 2.9 MHz (λ=517 μm). I_SP_ = 155 W/cm^2^, and a 100 ms pulse was repeated every 15 s for 12 trials. ***a***, Peak firing rate response from a single cell when the transducer was vertical (0°), showing strong modulation with a period of λ/2 as the transducer was moved away from the MEA. Distance is measured in terms of wavelength relative to a starting position, which was chosen to maximize the response. Also shown is the response when the transducer was tilted at an angle of 27° relative to vertical. ***b***, Raster plots and PSTHs of the response when the vertically oriented transducer was placed at the points indicated by colored circles in panel **a. c**, Left, Normalized population response for vertical (0°) and tilted (27°) transducers. Middle, FFT of firing rate vs distance, each point is an individual cell. The amplitude and phase are shown at a frequency of 2 cycles/λ for each cell that responds to ultrasound when the transducer is vertical (0°). In this plot, distance from the center represents the depth of modulation (log scale) of the response by the transducer position at a period of λ/2 and the angle is the phase of the 2 cycles/λ Fourier component, reflecting the transducer distance at which the response was maximal. The blue cross shows the mean population response. Right, FFT of firing rate vs distance when transducer was tilted at 27°.. **d**, Same as **c** except the 2.25 MHz transducer was operated at a carrier frequency of 1.9 MHz (λ=789 μm), and angle of the transducer when tilted was 21°, I_SP_ = 95 W/cm^2^. The maximal phase differs from **c** because the transducer was not repositioned to set the peak response at the starting position. For the tilted condition (21°), the distance traveled was only one full cycle (λ/2) with smaller step sizes.

### Potential contribution of radiation force to in vivo behavior

Although our findings indicate that higher acoustic frequency is more effective, consistent with radiation force, previous in vivo behavior experiments have shown that lower acoustic frequencies are more effective, (Ye et al., 2016). This would seemingly implicate mechanisms other than radiation force, and cavitation has been proposed as one potential mechanism. However, in previous experiments, because of diffraction, lower acoustic frequencies have always been accompanied by a large focal volume. As frequency is lowered, although radiation force would decrease with approximately the square of the acoustic frequency, the focal volume would increase with approximately the third power. Thus we considered whether radiation force could nonetheless quantitatively account for the in vivo results. We created a model that could predict the in vivo results with high correlation (r^2^=0.94). We found that summing radiation force across spatial volume by itself was not sufficient to explain the entire set of response curves (Fig. 8). We hypothesize that radiation force acting locally experiences an exponential saturating nonlinearity (see equation 1 in Methods and Fig 8a) such that a maximum strength of effect in a local region was reached at relatively low intensities for high frequencies (1.4 - 2.9 MHz); while low frequencies (0.4 – 0.5 MHz) do not saturate in the relevant range. Although higher acoustic frequencies would be more effective at activating a given local region, if this saturates at low intensities, and behavior is driven by the volume of activation, then the weaker but larger focal volume of lower frequencies recruits more of these regions and is thereby more effective at eliciting behavior. Following this saturating nonlinearity, activity was then spatially integrated across a volume whose parameters were fit. We found that optimal parameters for summation that matched the data was a depth of 1.2 mm and a radius of 3.7 mm. It is interesting that the depth parameter approximated the depth of mouse motor cortex, but the spatial radius parameter, if this model were correct, would imply that spatial integration occurred across an area larger than motor cortex. This model indicates that it is possible for radiation force to explain in vivo mouse data, but one must consider this model as an alternative hypothesis to be tested further.

**Figure 8.**
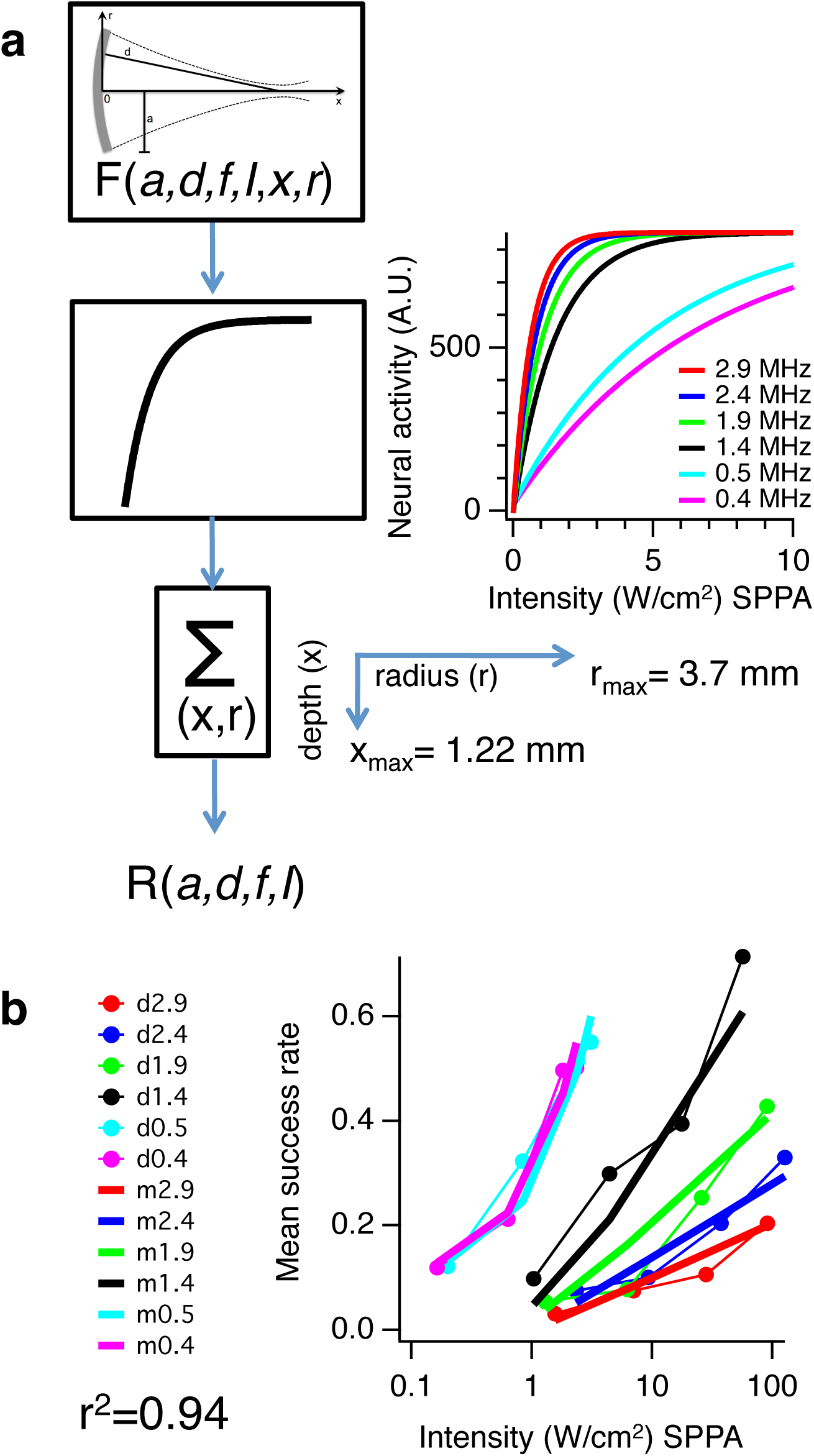
A radiation force model could explain in vivo brain data. **a**, Schematic of model that begins with an analytic expression used to calculate of radiation force in a cylindrical coordinate system based on transducer characteristics (*a* = radius of transducer, *d* = focal distance, *f*=frequency, *I* = intensity, x = axial distance, *r* = radial distance). This is the same model as in Fig. 6. At each point in space this radiation force is passed through a saturating non-linearity, so that radiation force produces a maximum localized effect. Plot at right shows that the relationship between neural activity at a single location and intensity saturates at higher but not low frequencies due to the increased radiation force at higher acoustic frequency. Neural activity is then summed over an optimized volume, with radius = 3.7 mm, depth = 1.22 mm to yield a single number representing mean success rate (% of time a stimulus results in behavioral muscle twitch). This is repeated for each different type of transducer, frequency, and intensity. **b**, Comparison of the model with in vivo mouse data. Thin lines connecting filled circles are data, thick lines are model, color of model matches color of data.

## Discussion

Our results show that ultrasonic neurostimulation in the retina produces radiation force and micron-scale displacement. A quantitative model of radiation force across multiple acoustic frequencies and power levels indicates that radiation force is the likely physical mechanism of action. We further show that standing waves can modulate neural activity, suggesting a potential new method to further control activity.

### Thermal effects of ultrasound

It is straightforward to estimate temperature rise produced by ultrasound based on I_SPTA_, duration of the pulse, density of brain tissue, specific heat capacity of brain tissue and the absorption coefficient (O’Brien, 2007). These estimates of temperature rise are very small (0.007°C, 0.04°C, 0.016°C respectively for (Lee et al., 2015; 2016a; 2016b)). Furthermore, these methods assume all energy goes to an increase in temperature, and do not account for heat loss by conduction or convection, so the actual temperature rise should be lower. This suggests that in these and other similar studies, ultrasound acts through mechanical and not thermal mechanisms.

We measured temperature rise under our experimental conditions using small (76 μm) thermocouples (J and K type, OMEGA) placed on the array with a retina pushed down on top of it. It is difficult to get accurate measurements because the rise in temperature is so small and localized. Thermocouples suffer from at least two sources of artifact. Ultrasound hitting the thermocouple will cause it to move relative to tissue and the resulting friction generates heat (known as the viscous heating artifact). However, in our measurements the thermocouple is attached to the bottom of the dish. The metal thermocouple will also conduct heat away. With perfusion, the temperature change is not measurable at 60 W/cm^2^ and 15 MHz, and without perfusion, we measure only 0.1-0.2 °C increase. For the 43 MHz transducer without perfusion we measured ~0.5°C increase at 30 W/cm^2^, well above the threshold for stimulation. (Menz et al., 2013). In a study using C. elegans (Kubanek et al., 2017), the authors measured behavioral response to ultrasound in wild types and mutants. They found that mutants lacking thermosensitive receptors behaved like wild type animals, while mutants that lack touch sensory neurons have an impaired response to ultrasound. They conclude that mechanical force, not heating, is the mechanism. Because we can only measure temperature changes when our perfusion is turned off and much higher power levels than the threshold for neural activity, we also conclude that the physical mechanism of action is mechanical, not thermal.

### Cavitation

In our study we found that higher acoustic frequencies were more effective than lower frequencies, thus ruling out cavitation as a possible physical mechanism. Using the same transducers, amplifier, frequencies and power settings that successfully stimulated in vivo mouse, we found that the lowest frequencies either did not stimulate retina cells (at 500 kHz, Fig 5c) or did so at much higher intensities compared to higher frequencies (1.9 MHz, Fig 5b). Cavitation can be measured with cavitation detectors which sense the subharmonics and harmonics produced by cavitation (Vykhodtseva et al., 1995; Gateau et al., 2011). To date, there is no study demonstrating the existence of cavitation using parameters for neurostimulation in the CNS. Cavitation requires gas bubbles; however, outside of the lungs and the digestive tract, biological tissue is generally bubble free (Church et al., 2008). Although is possible to pull a gas bubble out of solution; the pressures required are very high and tissue dependent (Nightingale et al., 2015). An in vivo sheep brain study with a 660 kHz carrier found that at least 12.7 MPa was required to measure a nucleation event with both passive and active cavitation detection (Gateau et al., 2011); whereas threshold pressures for low-power ultrasonic modulation in vivo brain studies are much less than 1 MPa (Naor et al., 2016). In another study using in vivo rabbit brain, the lowest intensity for the detection of cavitation was found to be 2000 W/cm^2^ at a carrier frequency of 0.936 MHz (Vykhodtseva et al., 1995); orders of magnitude higher than thresholds for ultrasonic neuromodulation for in vivo brain studies. The consequences of these high powered cavitation events are obvious damage to tissue (Vykhodtseva et al., 1995).

A hypothesis of ultrasonic neurostimulation is neuronal intramembrane cavitation excitation (NICE), which is a theoretical model that has been fit to empirical results (Krasovitski et al., 2011; Plaksin et al., 2014; 2016). The intramembrane cavitation hypothesis asserts that stable cavitation exists inside the cell membrane causing a change in cell capacitance that ultimately leads to action potential firing. This is a model of both a physical mechanism (cavitation) and a biophysical mechanism (change in membrane capacitance). Although this model has been fit to various in vivo experimental data, it does not describe our data because of the strong correlation of neural activity with acoustic frequency that we observe.

### Effects on ganglion cells

Previously we observed that blocking synaptic transmission with CdCl_2_ (Menz et al., 2013) abolished ultrasonic neurostimulation, indicating that we were not directly stimulating ganglion cells. One might assume, therefore, that the biophysical mechanisms of transduction are not present in the ganglion cell soma or dendrites. This could include specific types of ion channels, or properties of the presynaptic terminal. However, our present results show that little displacement was observed in the ganglion cell layer (Fig. 1), as the layer was close to the rigid MEA. Thus it may be that the ganglion cell soma can be directly activated by ultrasound if appropriate mechanical strain is applied. Further studies varying the geometry of the recording setup to produce mechanical strain at the ganglion cell level will be needed to assess whether ganglion cells can be activated directly.

### Potential Biophysical mechanisms

Leading candidates for biophysical mechanisms are mechanosensitive ion channels, capacitive effects from mechanical deformation of the cell membrane, and direct effects on endocytosis/exocytosis.

A simple biophysical mechanism that could transduce mechanical strain is a change in membrane capacitance, which can result from radiation force (Prieto et al., 2013). Alternatively, stretching, compressing or bending of the cell membrane may cause mechanosensitive ion channels to open or close. Mechanosensitive channels are found in all parts of the nervous system, serving different functions such as controlling osmotic pressure to guiding developing neurons (Orr et al., 2006; Haswell et al., 2011). Sensitive channels that are good candidates to convert mechanical stress from ultrasound into neural activity include Piezo, TRAAK, TREK-1, and TREK-2 (Brohawn, 2015; Syeda et al., 2016). In a study expressing mechanosensitive ion channels (two-pore-domain potassium family (K2P): TREK-1, TREK-2, TRAAK, and sodium channel NA_v_1.5) in *Xenopus* oocyte, ultrasound was found to significantly influence membrane current of the potassium channels and had a small effect on the sodium channel (Kubanek et al., 2016). In C. elegans, ultrasonic neurostimulation requires mechanosensitive channels (Kubanek et al., 2017).

It is known that very high static pressure will suppress synaptic activity. This is the physiological basis for High Pressure Neurological Syndrome (HPNS), a danger for deep-sea divers exposed to pressures greater than 1MPa (Jain, 1994; Aviner et al., 2016; Heinemann et al., 2017), but one from which divers fully recover without permanent damage. One potential mechanism of action is the abnormal functioning of calcium channels. An additional potential mechanism is a direct effect on exocytosis (Heinemann, Conti, Stühmer & Neher (1987). In general, multiple mechanisms of ultrasonic neurostimulation could operate under different conditions, including stimulus parameters or type of tissue.

### Relationship to in vivo studies

In the retina, higher carrier frequencies have lower thresholds and are more effective at generating neural stimulation, with effects predicted quantitatively by a radiation force model However, in vivo mouse studies using much lower frequencies (400 kHz-2.9 MHz) in which the output is muscle twitches (Ye et al., 2016), have shown that higher frequencies have greater thresholds. From these results, cavitation was proposed as a possible mechanism. Although different mechanisms may be involved in different systems, one must also consider that to date, lower acoustic frequencies have been applied with a focal spot of a larger volume due to effects of diffraction. The model we propose has a number of aspects that must be validated. The first is the localized saturating nonlinearity, which implies that either the biophysical mechanism such as ion channels, individual cells, or localized circuits reach a maximal effect of ultrasound. The second is the spatial scale of integration, including the radius and depth of the inferred effective area that drives behavior. Further measurements correlating displacement, neural activity and behavior will be needed to test this model.

Whether the exact details of the model we propose are correct, one must consider that even though lower acoustic frequencies may be less effective at creating radiation force, effects that are integrated over a volume larger than the focal spot will increase with the third power of the frequency, such that overall lower frequencies may be more effective even though per unit volume lower frequency is less effective. In the present studies, the retina is effectively twodimensional with respect to the frequencies tested. Furthermore, acoustic frequencies that we used of 15 MHz and lower have spot sizes larger than the ganglion cell receptive field center. Thus, the varying spot size may have less of an effect in the retina such that only the frequency dependence of radiation force plays a role. Further studies controlling for spot size in vivo will be needed to assess this relative effects of radiation force and focal volume.

### Pressure phosphenes

It has been known since ancient Greece that mechanical deformation of the eyeball generates pressure phosphenes (the appearance of light when there is none). Although it is still not known which cells in the retina are responsible, this is a clear demonstration that mechanical strain can result in ganglion cell activity. In studies with deformation of cat eyeball combined with electrical recordings from single optic tract axons it was shown that different ganglion cells respond differently; i.e., On–center and Off-center responded to ultrasound with a polarity consisted with visual stimulation (Grusser et al., 1989a; 1989b). To account for this antagonism, ganglion cells were likely not being directly stimulated, and it was proposed that other cells in the network were being stimulated, likely in the outer retina. Most importantly they conclude that mechanical strain is the cause, not retinal ischemia from high intra-ocular pressure. The authors speculate that horizontal cells might be the most sensitive to stretching of the retinal surface area since they lie laterally in the retina. A phenomenon that has been known for thousands of years supports the concept of mechanical strain on neurons as the cause for this neural stimulation.

## Conclusion

There exists a strong theoretical and empirical understanding of using radiation force and standing waves to exert mechanical effects in the fields of acousto-fluidics (Bruus, 2012; Lenshof et al., 2012) and elasticity imaging (Doherty et al., 2013). Here we show a new application of these principles to ultrasonic neurostimulation. Our findings suggest that future approaches of ultrasonic neurostimulation should maximize radiation force induced mechanical strain while minimizing power. An understanding of the physical mechanism of action will allow studies in this area to pursue how radiation force might be manipulated to optimize stimulation and simultaneously provide insights into biophysical mechanisms.

---

**Supplemental Figure 1.**
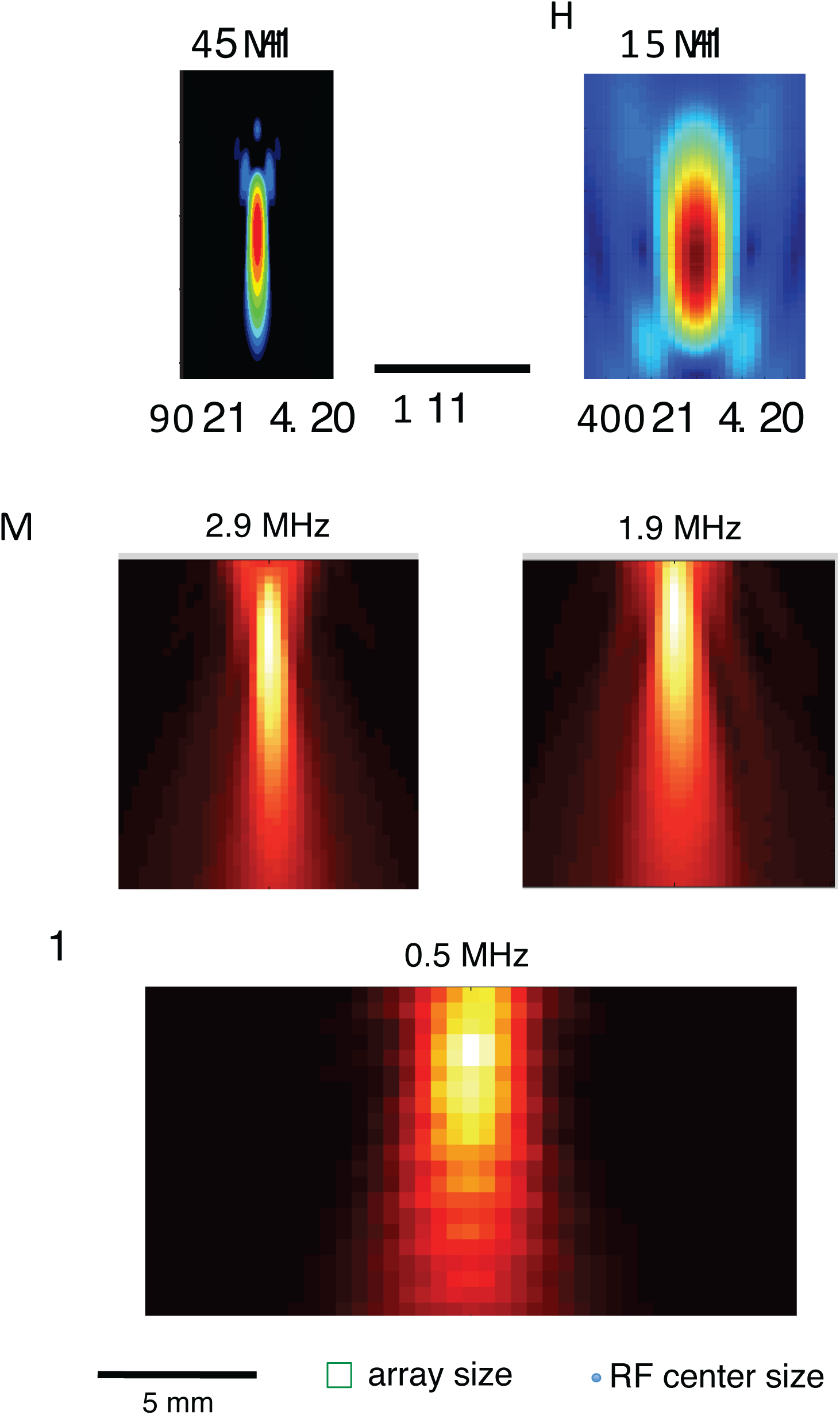
Spatial distribution of intensity for different transducers and carrier frequencies. **a**, XZ plot of intensity for 43 MHz transducer, having a 90 μm lateral and 1330 μm longitudinal focal volume (-3 dB). **b**, 15 MHz transducer. 1 mm scale bar applies to **a** and **b. c**, A 2.25 MHz transducer was operated at two different frequencies. Left, 2.9 MHz and Right 1.9 MHz. **d**, 0.5 MHz transducer. 5 mm scale bar applies to **c** and **d**. Bottom, array size and typical ganglion cell RF center size.

## Acknowledgements

We thank M. Maduke, M. Prieto, J. Kubanek, D. Palanker and J. Brown for helpful discussions. This work was supported by a grant from the NIBIB and the Stanford Neurosciences Institute.

## Author Contributions

M.M., P.Y., K.B.P, P.K-Y. and S.A.B. designed the study, M.M. and P.Y. performed the experiments, M.M. analyzed the data. K.F. performed simulations, and M.M. and S.A.B. wrote the manuscript.

## Methods

### Electrophysiology

Multielectrode array (MEA) recordings were performed as described (Manu & Baccus, 2011) The isolated retina of the tiger salamander was adhered by surface tension to a dialysis membrane (Spectrapor 7 50000, Fisher Scientific) attached to a custom Delrin holder. The holder was placed on a motorized micromanipulator (MP-385-2, Sutter) and lowered onto a multielectrode electrode array (ThinMEA, Multichannel Systems) ganglion cell side down. For 43 MHz experiments where the focal spot < 100 μm, a high density array was used (5x6, 10 μm diameter electrode, 30 μm spacing). For all other lower frequency experiments, a lower density array was used (8x8, 10 μm dia., 100 μm spacing), which better matches the focal spot size. Full field flashes from a red LED were sometimes used to verify that ganglion cells were responding normally to visual stimuli, especially if conditions of ultrasound stimulation did not show a response. Error bars are SEM unless otherwise noted.

### Ultrasound transducers and stimuli

We used four different transducers, 43 MHz (custom), 15 MHz (Panametrics, A319S, 0.5” dia, 2” FL), 2.25 MHz (Olympus, V305) and 0.5 MHz (Olympus, V301) in order to span a large frequency range. The 2.25 MHz transducer has a relatively wide bandwidth and was operated at multiple frequencies (1.9 and 2.9 MHz). Transducers (15, 2.25 and 0.5 MHz) were fitted with a water–filled cone and mounted on a motorized micromanipulator (MP-385-2, Sutter). A camera from below was used to position the transducers so that the center of the focal spot was in the center of the array. Transducers were lowered into the bath above the retina, and height was adjusted so that the focal point was on the retina. Ultrasound propagated from the transducer, through the water-filled cone, perfusion fluid, dialysis membrane, retina, and then reflected off the glass/metal surface of the MEA (Fig 8). A function generator (model 8116A, Hewlett-Packard) provided the carrier frequency that was gated by the analog output of a National Instruments DAQ board. This signal was amplified by a 50 dB RF power amplifier (model 320L, Electronic Navigation Industries) and fed into the transducer. A hydrophone was used to measure power output from the water–filled cones into a tank of water as a function of three spatial dimensions (Fig 9), except for 43MHz, which is too high for a conventional hydrophone, and for which power was extrapolated from hydrophone measurements at 20 MHz. All power measurements in this paper are the spatial peak, I_SP_ because with a 100% duty cycle (continuous wave) I_SPPA_ = I_SPTA_ (i.e., pulse average = temporal average) in free space (water tank). These free space hydrophone measurements are not corrected for the reflection off of the MEA under experimental conditions and the resulting standing wave. The free space measurements we have provided are useful for reproducing our results and making relative comparisons across carrier frequencies. However, these measurements do not accurately describe the actual power distributed in space under experimental conditions where we have standing waves between the transducer and the MEA. Continuous wave is used for all experiments so the only relevant parameters are carrier frequency, power, pulse duration, and repetition rate, which are given for each experiment.

### Imaging

The styryl dye FM4-64 was bath applied by immersing the isolated retina in a concentration of 82 μM (100 μg in 2 ml) FM4-64 in oxygenated Ringer’s for one hour prior to placement on the MEA. This dye inserts itself in the outer leaflet of the cell membrane where it becomes fluorescent, allowing us to image changes in position and shape of the cell membrane with ultrasonic stimulation.

A custom two-photon laser scanning microscope in the inverted configuration was used to image the retina during ultrasonic stimulation. A simplified diagram is shown in Fig 8a. Excitation at 970 nm from a Ti:sapphire laser (Tsunami, Spectra-Physics) was focused on the retina by a x 40 1.2 NA (Zeiss) objective and the epifluorescence passed through an emission filter (FF01- 725/150-25) and laser-light blocking filter (Semrock, FF01-680/SP-25), which was then collected by a PMT (H7422-P, Hamamatsu). The imaged area was selected to cover the ultrasound focal spot. We recorded a frame of 512x128 pixels at a rate of 18.6 frames per second for 1000 frames at one level in the retina. We stared at the MEA and collected images in one μm steps for a total of 120 microns, covering the entire retina in depth, imaging 1000 frames at every step. The average laser power was set to 10mW. ScanImage (now supported by Vidrio Technologies) software was used to record images.

The mirror position of the scanning galvanometers was recorded on the same computer that generates and records the ultrasound stimuli (one second on, one second off, 40W/cm^2^). This allowed us know the timing of the image at any pixel relative to the ultrasound stimulus. The laser scanning and ultrasound stimulus are not synchronized, such that any given pixel will be recorded at random times during the two second period of ultrasound stimulation. In theory we can get arbitrarily high temporal resolution in this way with a sufficiently large data set; in practice we binned the data in 10 ms bins for a sufficient signal-to-noise ratio. To compute vector fields reflecting the effect of ultrasound, we used unwarpJ, which is an imageJ plug-in that performs spline-based elastic registration of two images. We compared steady state images in the ultrasound on and off conditions.

Because of the large area of scanning at a high frame rate (20 Hz), we corrected for most distortion at the edge of the frame by computing the average actual mirror position based on control experiments recording mirror positions with slow mirror velocities and no distortion and then recording actual mirror positions at high velocities. This distortion does not affect our analysis, which is based on changes in the images as a function of time in the cycle of ultrasonic stimulation.

### Modelling radiation force to explain retinal displacement

To model the observed displacement as an effect of radiation force, we assumed the ultrasound field at 43 MHz was transmitted through a multilayered medium composed of water, retina, glass, and air, and calculated assuming 40 W/cm^2^ incident power. The retina layer was 100 um thick and the glass layer was 180 um thick. Water and air media were assumed to be half-spaces. We then calculated the radiation pressure on the retina-water and retina-glass interfaces (Lee and Wang, 1993). The density of the retina was set to 1000 kg/m^3^. The speed of sound in the retina was estimated based on the reflection coefficient at the water-retina interface. Lab measurements indicated a reflection of +0.2 at the water-retina interface. Estimated radiation pressures were then used in COMSOL Multiphysics finite element software to calculate the deformation of the retina in response to ultrasound. The retina was considered to be an incompressible material (i.e., Poisson ratio of 0.5). We determined the value of the elastic modulus (2.3 kPa) which gives 4 μm of displacement as seen in the data. This estimate of the elastic modulus necessary to obtain the desired displacement is close to the measured range found in the inner retina (0.94 - 1.8 kPa) with the scanning force microscopy method (Franze et al., 2011). Since soft tissues such as retina exhibit large deformation with nonlinear strains, a large deformation model was used to estimate the displacement field in the retina.

### Modelling radiation force to explain neural activity

A quantitative radiation force model was used to fit population activity in the retina. The model is based on analytic equations valid for linear low-amplitude ultrasound in free space (Eqs 13,14,16 in (Rudenko et al., 1996) and Eq 9 from (Ye et al., 2016)). We did not account for the reflection off of the MEA and the resulting standing waves. For 1.9 MHz, where standing wave effects are large, we use the data from the transducer position with the lowest threshold. The analytic expression takes as input the absorption coefficient (retina is similar to brain, so we used the same parameters as (Ye et al., 2016)), the carrier frequency (*f*), intensity (*I*), radius of the transducer (*a*), and focal length (*d*), to estimate radiation force in three-dimensional space expressed in a cylindrical coordinate system where x is axial distance from the transducer and *r* is the radial distance away from the central axis. To fit the normalized population neural response in the retina (Fig 5ab) to the model, the maximum radiation force was calculated for each intensity and transducer combination and passed through a sigmoid:

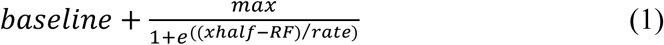

The sigmoid represents neural nonlinearities where “max” is the saturation level (0.73), “xhalf” is the x value at half-max (0.5), “rate” is the gain (0.25), “baseline” determines the output at zero RF, and RF is the maximum radiation force generated at a given intensity for a given frequency. Higher frequencies require less intensity to generate the same radiation force. There were four free parameters in the model, all describing the sigmoidal output non-linearity. The four free parameters were found by minimizing the total rms error between the data and the model.

### Radiation force model of *in vivo* behavior

To fit the mouse data to a radiation force model (Fig. 7) we used the same analytic equations as for the retina model to compute radiation force in 3-dimensional space for each transducer frequency and intensity. As with the retinal model, we did not account for standing waves. Unlike the retinal model, the conversion from radiation force to neural activity is performed at each spatial location with a non-linear function, the exact form of the non-linearity is not critical, but the threshold should be low and it should saturate (Fig. 5b), we used the following expression:

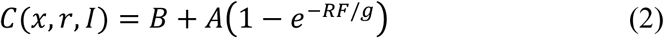

where *C* is the non-linear localized response at location (*x,r*), *B* is the baseline response at the lowest intensity tested, *g* is the gain, *A* will determine the saturating response, and *RF* is the radiation force. The effect of this non-linearity at different frequencies is shown in Fig 8a. We allowed the parameter *B* to take on a different value for the two lowest frequencies (400 kHz and 500 kHz) because their response at minimum intensity is higher than the other frequencies (Fig 7c). It is possible that higher acoustic frequencies are actually inhibiting spontaneous activity at low intensities (see Fig 5, high frequencies – focused, from (Ye et al., 2016)). The non-linearity is interpreted as a local biophysical mechanism (such as mechanosensitive ion channels in individual cells) that has a very limited dynamic range. The final component of the mouse brain model is to sum all of these local responses *C* over a volume specified by the parameters *r_max_* and *X_max_* which are free parameters. This summation represents the hypothesis that circuits responsible for behavior could be summing activity over a large volume. The model has a total of five free parameters, three associated with the non-linearity and the two spatial parameters.

